# Boosting Hyperalignment Performance with Age-specific Templates

**DOI:** 10.1101/2025.02.19.639148

**Authors:** Yuqi Zhang, Maria Ida Gobbini, James V. Haxby, Ma Feilong

**Affiliations:** Dartmouth College; University of Bologna; University of South Carolina

**Keywords:** fMRI, hyperalignment, brain aging, individual differences, functional connectivity

## Abstract

Hyperalignment aligns individual brain activity and functional connectivity patterns to a common, high-dimensional model space, resolving idiosyncrasies in functional–anatomical correspondence and revealing shared information encoded in fine-grained spatial patterns. Given that the brain undergoes significant developmental and functional changes over the lifespan, it is likely that certain features in brain functional organization are more prominent in certain age groups than others. In this study, we examined whether age-specific functional templates, as compared to a canonical template, could enhance alignment accuracy across diverse age groups. We used the Cambridge Centre for Ageing and Neuroscience (Cam-CAN) dataset (18 to 87 yo) to build age-specific templates and tested their performance for analyzing data in young and old brains in both the Cam-CAN dataset and the Dallas Lifespan Brain Study (DLBS) dataset (20 to 90 yo). We found the congruent age-specific template outperforms the incongruent template for various analyses, including inter-subject correlation of hyperaligned connectivity profiles and predicting individualized connectomes and brain responses to the movie using the template. The results are consistent across both datasets. This work enhances our understanding of age-related differences in brain function, highlights the benefits of creating age-specific templates to refine hyperalignment model performance, and may contribute to the development of age-sensitive diagnostic tools and interventions for neurological disorders.

## Introduction

Information encoded in the cortex can be decoded from fine-grained patterns of cortical activity via multivariate pattern analysis of fMRI data (Haxby et al., 2001, 2014). However, there is considerable variability across individual brains when encoding the same information (Cox & Savoy, 2003). Traditional approaches to brain alignment often fail to capture the fine-grained functional correspondence between brains due to topographic variability. Hyperalignment is a computational approach for modeling how the brain encodes shared information across individuals, despite individual variability in cortical topographies (Haxby et al., 2011, 2020). Hyperalignment can be trained using either brain responses (Haxby et al., 2011; Guntupalli et al., 2016) or functional connectivity (Guntupalli et al., 2018), and it captures both shared coarse- and fine-scale information encoded in the brain. This approach also enhances the reliability of individual differences by affording analysis of differences in the fine-grained structure of the functional connectome (Feilong et al., 2018, 2021).

Given that the brain undergoes substantial developmental, structural, and functional changes across the lifespan, brain functional organization may demonstrate age-specific features that are not prominent in other age groups. These age-related variations can impact the alignment of neural data across individuals. Consequently, the performance of hyperalignment models in human connectomics may be significantly influenced by the age of the individuals used to build the templates. In this study, we investigate whether the incorporation of age-specific templates enhances the performance of hyperalignment models. Examining this aspect could not only improve hyperalignment accuracy across diverse age groups but also inform age-sensitive intervention strategies. Individual differences in the brain arise from both aging and neurological diseases and brain injuries. This knowledge also could open new pathways for creating more effective diagnostic tools for neurological disorders (Anderson et al., 2021, 2024).

We investigated this problem by developing age-specific hyperalignment templates using the Cambridge Centre for Ageing and Neuroscience (Cam-CAN) dataset and evaluating their performance with three indices: (a) inter-subject correlation (ISC) of connectomes, (b) prediction accuracy of individualized connectomes, and (c) prediction accuracy of individualized brain responses to the movie. Across all three analyses, we found consistent advantages of congruent age-specific templates (i.e., those constructed using data from the same age group) over incongruent templates. Together, these results demonstrate the importance of accounting for age-specific features of brain functional organization when applying functional alignment methods.

## Methods

### fMRI dataset

The Cambridge Centre for Ageing and Neuroscience (Cam-CAN) dataset comprises fMRI data of over 600 people from a cross-sectional adult lifespan (18 to 87 years old) population-based sample, with approximately 25 minutes of fMRI data per individual (Taylor et al., 2017). All MRI datasets were collected at MRC-CBSU using a 3T Siemens TIM Trio Scanner with a 32-channel head coil, 3.0 × 3.0 × 4.44 mm^3^ voxels and 20% gap. Each individual’s fMRI data were collected during three different tasks:

1. 8 minutes 40 seconds of resting state with eyes closed (‘rest’);
2. 8 minutes of movie-watching of the film “Bang! You’re dead” (‘bang’);
3. 8 minutes 40 seconds of a sensorimotor task (‘smt’) during which participants were asked to press a button upon the presentation of a visual and/or auditory stimulus.

Both resting state and the sensorimotor task used single echo sequences with TR = 1970 ms, TE = 30 ms, flip angle = 78°, and the movie-watching task used multi-echo sequences with TR = 2470 ms, TEs = 9.4, 21.2, 33, 45, 57 ms, flip angle = 78°. 646 participants have the full records of all three functional scans (age distribution shown in Table 1).

**Table 1.** Age group division and corresponding training/test set sizes.

| Age Group | Age Range | Number of Participants | Training Group | Test Group |
| --- | --- | --- | --- | --- |
| Young | 18–45 | 215 | 144 | 71 |
| Old | 65–90 | 216 | 144 | 72 |

We classified the participants into three different age groups—young, mid, and old—with each group consisting of approximately the same number of individuals. Our analyses focus on the young and old groups.

### Preprocessing

We preprocessed all MRI data using fMRIPrep (Esteban et al., 2019), with version 20.2.7. For each participant, the cortical surface was reconstructed using the high-resolution structural scans. The functional data were corrected for head motion, projected onto the cortical surface model, and resampled to the onavg-ico32 cortical surface template, which uniformly samples different parts of the cortex (Feilong et al., 2024). After that, we used linear regression to partial out nuisance regressors from our data. The list of nuisance regressors includes 6 head motion parameters and their derivatives, framewise displacement, global signal, 6 aCompCor components from cerebrospinal fluid and white matter, and polynomial trends up to the 2nd order. After the regression, we normalized the time series of each cortical vertex to zero mean and unit variance.

### Individual connectome calculation

We computed two kinds of connectomes based on different functional scans:

1. using both resting state and sensorimotor task data,
2. using movie-watching data,

This division allows us to examine template performance using both functional connectivity and response time series to the movie. To calculate functional connectivity, we downsampled the data matrices for each individual and each task from ‘onavg-ico32’ space (19341 vertices) to ‘onavg-ico8’ space (1210 vertices) and used the time-series for downsampled data as connectivity targets. Correlations between connectivity target time-series and vertex time-series are indices of functional connectivity. Each vertex’s connectivity profile is a vector of 1210 elements, where each element is the correlation between the time series of the vertex and the time series of a connectivity target. We z-scored the connectivity profile for each of the 19341 vertices, and these connectivity profiles collectively form the connectome matrix of the individual and task.

### Hyperalignment template creation

For each age group (young and old), we built the templates using approximately two thirds of the participants and withheld the remaining one third for testing. Given there are multiple ways to choose the participants, we repeated the procedure 10 times, each time randomly choosing two thirds of the participants without replacement. The participants used for evaluation were always independent from the participants used for training.

We combined fMRI data of different tasks to maximize the amount of data we use to compute the connectomes, given that differences in connectomes based on different tasks are much smaller in scale compared to individual differences (Gratton et al., 2018). Each whole brain template was created using a searchlight-based algorithm (Feilong et al., 2023, section 4.2) with a 20 mm searchlight radius, based on the training group participants’ connectome data calculated from the resting state data and sensorimotor task data. There are 19341 overlapping searchlights, each containing an average of approximately 121 vertices (range: 44 to 187). For each searchlight, we created a local template so that its representational geometry and topographies are representative of the training participants. We first concatenated the vertices of all participants and performed a PCA on the concatenated data, to derive a PC template that reflects the representational geometry of the searchlight, where *M*_(PC)_ is the PC template,*B*_(*p*)_ is the local data matrix of the *p*-th participant, and *n*is the number of participants.

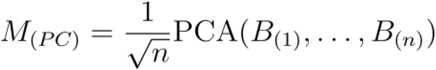

We then applied a rotational matrix *R* to the PC template to minimize the topographic differences without changing the information content, where ‖ · ‖_*F*_ is the Frobenius norm.

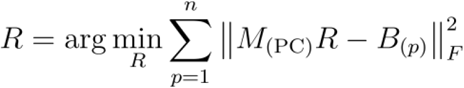

To find the solution *R*, we applied the orthogonal Procrustes algorithm to concatenated data matrices (see formula below).

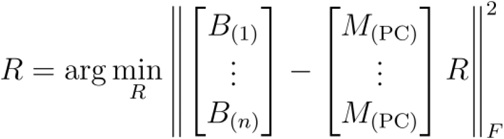

The local templates *M*_(PC)_*R* searchlights were then combined to form the whole-brain template. For each participant *p*, we derived two transformation matrices using Procrustes algorithm: *R*_(*p*)_, which projects data from the participant’s native cortical space to the template space and was used in inter-subject correlation (ISC) analyses, and 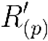, which projects data from the template space back to the participant’s native cortical space and was used in the prediction analyses (Figure 1, see also Supplementary Figure 1). Note that these matrices differ from the rotation matrix *R* used to derive the local template.

**Figure 1.**
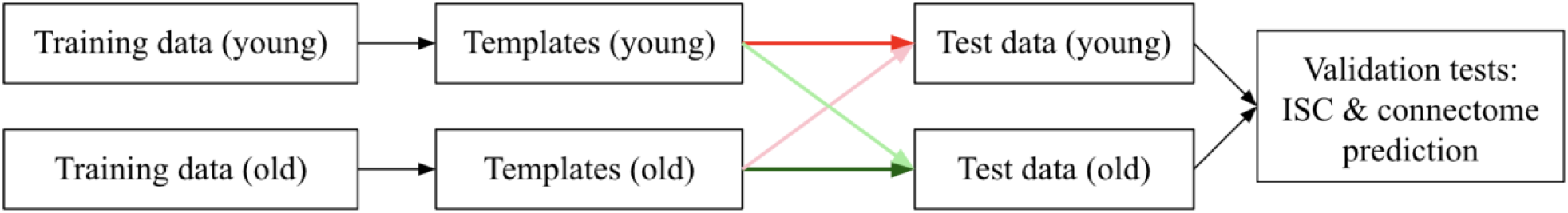
Schematic of the procedure for building and testing hyperalignment templates. Darker arrows indicate congruent templates (i.e., from the same age group), whereas light arrows indicate incongruent templates.

Given that functional connectomes may differ across tasks, for each template based on resting state and sensorimotor data, we also computed a corresponding template based on the movie-viewing data. This was used in the analysis of predicting individual connectomes, as described below.

### Inter-subject correlation (ISC)

We computed the inter-subject correlation (ISC) of connectivity profiles in the common model space to evaluate template performance, where higher ISCs indicate better performance. For each participant in the test set, we computed the participant’s hyperalignment transformation matrix *R*_(*p*)_ using the connectome based on resting state and sensorimotor task data, and applied the transformation matrix to the original movie-watching responses to compute the hyperaligned connectome. This ensures that the data used to compute hyperalignment transformation are independent from the data used to evaluate performance. We computed ISC as the correlation between the participant’s connectivity profile and the average profile across other test participants in the same age group (Figure 1). Each participant has up to 20 ISC maps (10 young templates and 10 old templates), and we averaged ISC maps of the same kind (young or old) for each participant. We converted *r* values to *z* values (Fisher transform, arctanh(r)), averaged the *z* values, and converted them back to *r* values. Based on the analysis, we either averaged them across cortical vertices (Figures 2a, 2b, Supplementary 3a, 3b) or participants of the same age group (Figures 2c, Supplementary 3c).

**Figure 2.**
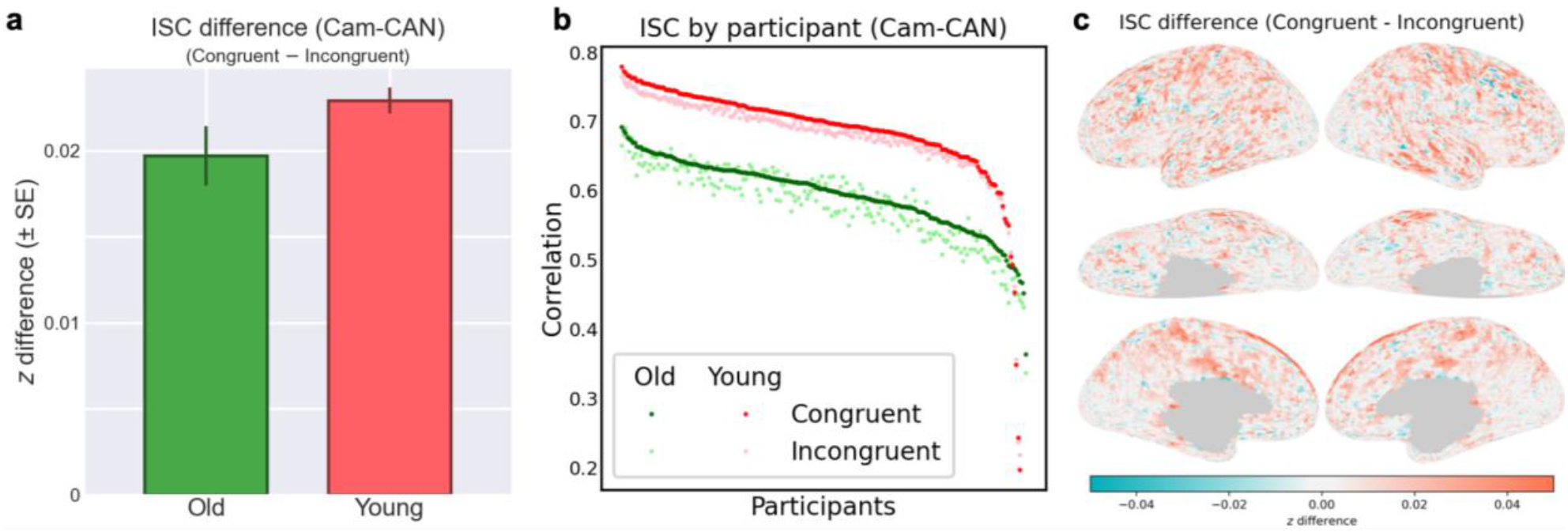
Inter-subject Correlation Results (Cam-CAN). (a) The average *z* (± SD) for the old group was 0.6897 ± 0.0759 for congruent templates and 0.6700 ± 0.0839 for incongruent templates. The mean ISC difference (± SE) was 0.0197 ± 0.0017, *t*(212) = 11.33, Cohen’s *d* = 0.7766 and *p* < 10^-22^. The average *z* (± SD) for the young group was 0.8566 ± 0.1151 for congruent templates, and 0.8337 ± 0.1069 for incongruent templates. The mean ISC difference (± SE) was 0.0229 ± 0.0008, *t*(209) = 30.09, Cohen’s *d*= 2.0765 and *p* < 10^-77^. (b) Scatter plot of each participant’s average ISC value derived from different congruent and incongruent templates. (c) Topographic distribution of ISC difference across different brain regions.

### Predicting individual connectome

We also tested the prediction of connectomes in held-out test participants’ native anatomical space (Figure 1). We derived each participant’s predicted movie-viewing connectome as the matrix multiplication of the movie-viewing template and the individual’s transformation matrix 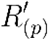, derived from resting state and sensorimotor task data. We then compared the predicted connectome with the participant’s actual movie-viewing connectome based on the correlation of the corresponding connectivity profiles in these two connectomes.

### Predicting individual responses to the movie

Similarly, we predicted individual responses to the movie using the same templates created for the congruent and incongruent age groups. Using the Individualized Neural Tuning (INT) model (Feilong et al., 2023), each individual’s connectome was modeled as the same template connectome with an individualized transformation, and we established correspondence between the modeled connectome and the brain responses to the movie using the same group of training participants used to build the template. In other words, we applied the INT model to enrich the template so that it could predict individualized brain responses in addition to individualized connectomes based on the template. We then evaluated the template’s performance using correlation coefficients between the measured response time series to the movie and the predicted time series.

### Validation dataset

The results based on the Cam-CAN dataset demonstrate that hyperalignment templates from the same age group work better than templates from age-incongruent templates, and this effect generalizes across tasks (from resting state and sensorimotor to movie-watching) and scan protocols (from single-echo sequences to multi-echo sequences). We further tested the generalizability of our results using an independent dataset—the DLBS dataset (Park et al., 2025), which was collected with a different scanner, a different group of participants, and different tasks (See Supplementary).

## Results

We evaluated the performance of the templates using three metrics: (a) ISC of functional connectivity in the common template space, where higher ISCs indicate better alignment of connectivity in the template space; (b) similarity between model-predicted and measured connectomes in each participant’s native anatomical space; and (c) similarity between model-predicted and measured neural responses to the movie, where higher similarity suggests the template contains more relevant information and better predicts individualized connectomes and movie responses. Across all metrics, we found that congruent age-specific templates outperformed age-incongruent templates.

### Inter-subject correlation

We first evaluated the performance of templates constructed from both young and old age groups on ISCs of connectomes in the common model template space. We computed the z-difference (difference in Fisher z transformed correlations) between post-alignment ISC values derived from congruent and incongruent templates (Figure 2a). Hyperalignment based on congruent age-specific templates yields higher ISCs in both age groups, with a more pronounced effect in the young group. Breaking down the data by participant, we found that most participants (97.6% in the young group, 74.2% in the old group) exhibited higher ISCs when comparing the results for congruent and incongruent templates (Figure 2b). We also evaluated the ISC performance of an intermediate middle-aged cohort using a middle-aged template; these results are provided in Supplementary Figure 4. The topography of mean ISC differences displayed in Figure 2c illustrate that congruent templates perform better in the frontal, temporal, and parietal lobes. The ISC for the young group was consistently higher than that for the old group for both congruent and incongruent templates. This reflects, at least in part, differences in data quality between the two age groups (Supplementary Figure 2).

### Predicting connectomes

Higher ISCs can arise from both (a) better alignment across participants and (b) filtering out noise during hyperalignment transformations. Though it is unlikely that our ISC results were driven by reduced noise, to rule out the possibility, we assessed how well different templates predict each participant’s connectome.

For each individual, we calculated the correlation between the predicted connectome, generated using young and old training group hyperalignment templates calculated with rest and smt fMRI data, and the actual connectome, calculated using movie watching fMRI data. A higher correlation indicates a more accurate prediction, which means better performance of the hyperalignment template.

We compared the performance of congruent templates (built from the same age group) and incongruent templates (built from a different age group). For each participant in the test sets of both age groups, we calculated the predicted connectome using 20 templates—10 from each age group. We then computed the correlation for each participant and averaged the individual results across templates in the same age group. Figure 3a shows the z-difference between correlations calculated using congruent and incongruent templates for each participant in both groups. The results clearly indicate that predictions using congruent templates are more accurate than those using incongruent templates. Examining the participant breakdown in Figure 3b, we observed that almost all participants (98.6% in the young group and 94.4% in the old group) have a better connectome prediction using congruent templates than using incongruent templates from the other group. Additional results predicting the fine-grained connectome for the middle-aged cohort using the middle-aged template are detailed in Supplementary Figure 5. We also evaluated connectome prediction accuracies using templates constructed from 10-year age increments. The results show a continuous gradient of age-related divergence (Supplementary Figure 6). When predicting data for the 80–90 cohort, the 20–30 template performs the worst and the performance steadily improves as the template age gets closer to the target demographic. This systematic gradient further supports our main finding: the penalty for using an incongruent template increases with the discrepancy between the template age and participant age. The topographic distribution of mean differences between the predicted connectomes using congruent and incongruent templates are displayed in Figure 3c. Congruent templates generally perform better in the frontal and parietal lobes—regions primarily responsible for cognitive functions—which can be significantly influenced by age.

**Figure 3.**
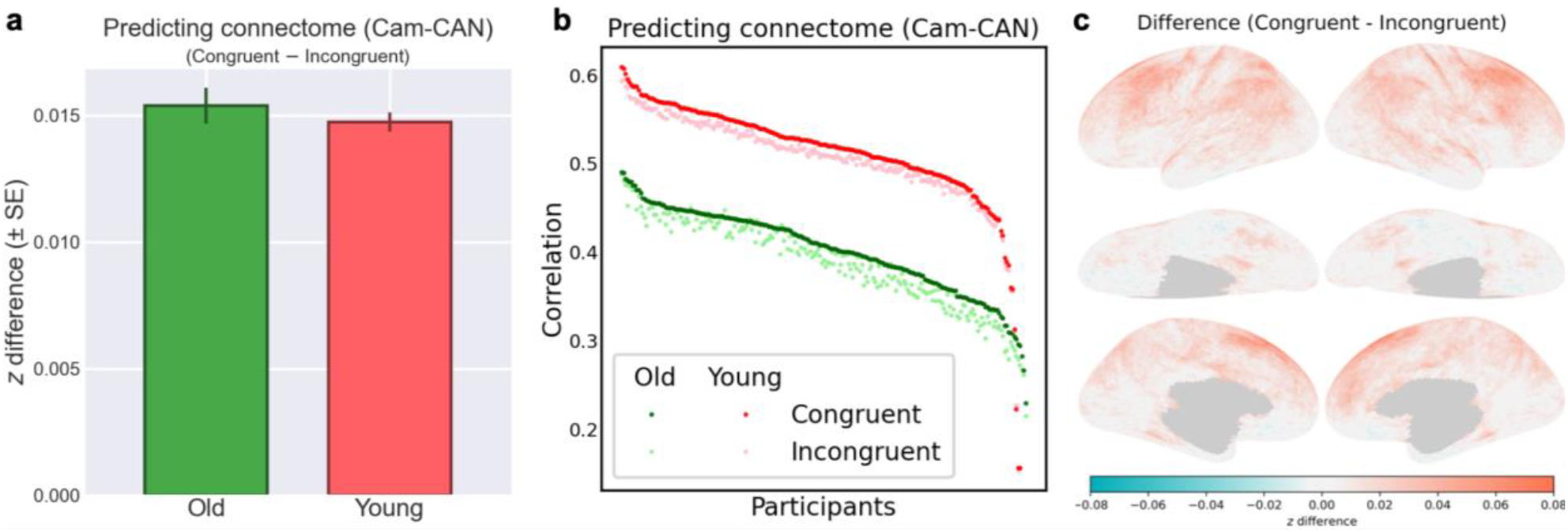
Prediction performance comparison between congruent and incongruent templates. (a) The average *z* (± SD) for the old group was 0.4279 ± 0.0558 for congruent templates, and 0.4126 ± 0.0597 for incongruent templates. The mean difference of predicted connectome (± SE) was 0.0154 ± 0.0007, *t*(212) = 21.91, Cohen’s *d* = 1.5014 and *p* < 10^-55^. The average *z* (± SD) for the young group was 0.5716 ± 0.0779 for congruent templates, and 0.5568 ± 0.0743 for incongruent templates. The mean difference of predicted connectome (± SE) was 0.0147 ± 0.0004, *t*(209) = 39.07, Cohen’s *d* = 2.6963 and *p* < 10^-97^.(b) Scatter plot of each participant’s average correlation value derived from different congruent and incongruent templates. (c) Topographic correlation difference across different brain regions.

To evaluate the influence of age on the performance of hyperalignment template across the entire lifespan, we computed the correlation between actual connectome and predicted connectome of all participants using both young templates and old templates. As an individual’s age becomes more distant from the template age group, the relative performance of the template decreases (Figure 4a, 4b).

**Figure 4.**
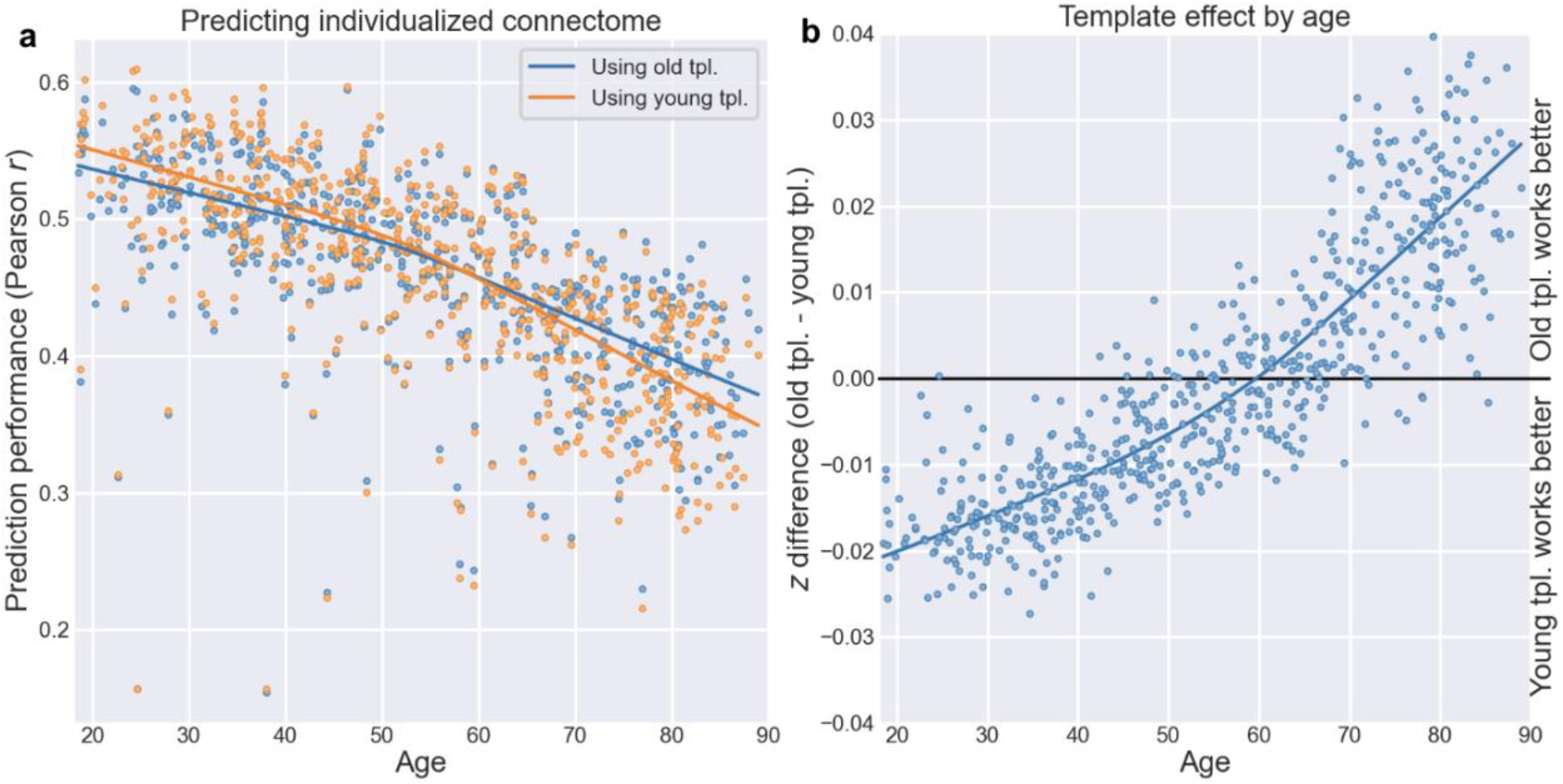
Comparison of individual connectome prediction across all age spans. (a) Scatter plot showing Pearson correlation between actual connectome and predicted connectome derived from young and old templates for individuals across all age spans (b) Correlation difference among individuals between two age group templates, ordered by age.

Overall, congruent age-specific templates better predict participants’ connectomes than incongruent templates. As the participant’s age diverges further from the template age group, the prediction accuracy gets worse. This result indicates that age independent templates are necessary for improving the performance of hyperalignment.

### Predicting brain responses to the movie

Besides functional connectivity, congruent templates also performed better than incongruent templates in predicting neural responses to the movie (Figure 5). These results indicate that the more congruent the template age group is with a participant’s age, the more accurate the prediction becomes.

**Figure 5.**
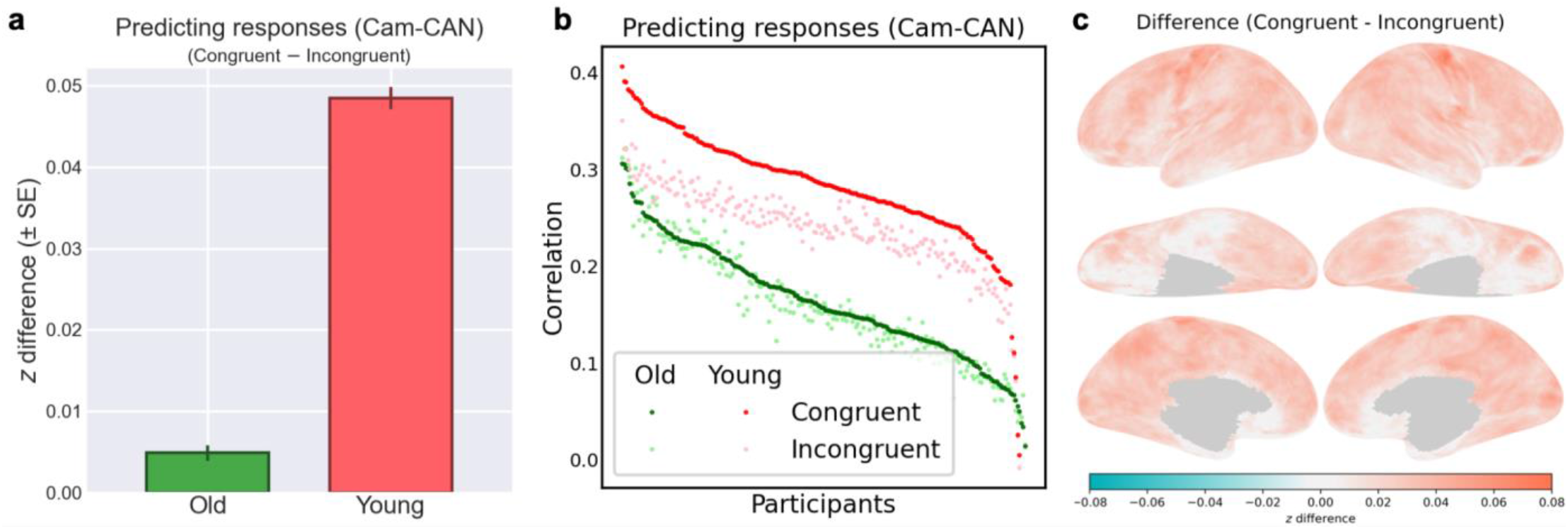
Predicting brain responses to the movie based on congruent and incongruent templates. (a) The average *z* (± SD) for the old group was 0.1661 ± 0.0600 for congruent templates, and 0.1612 ± 0.0636 for incongruent templates. The mean difference (± SE) was 0.0049 ± 0.0010, *t*(212) = 5.02, Cohen’s *d* = 0.3443 and *p* < 10^-5^. The average *z* (± SD) for the young group was 0.2925 ± 0.0627 for congruent templates, and 0.2441 ± 0.0490 for incongruent templates. The mean difference (± SE) was 0.0485 ± 0.0013, *t*(209) = 35.99, Cohen’s *d* = 2.4833 and *p* < 10^-90^. (b) Scatter plot of each participant’s average predicted response values derived from different congruent and incongruent templates. (c) Topographic distribution of predicted response difference across different brain regions.

We used the same 10 functional templates for each age group, based on different participants randomly chosen as described earlier, to calculate individual transformation matrices and predict movie responses. We then focused on differences in performance between using congruent and incongruent templates.

Next, we computed correlation coefficients between the measured response time series to the movie and the predicted time series to evaluate template performance. The correlation coefficients were Fisher-transformed to *z* values and aggregated either across different parts of the brain (Figures 5a, 5b) or across participants (Figure 5c). Prediction performance for both groups was better when using congruent templates than incongruent templates, with the young group showing a larger congruency effect than the old group.

## Discussion

In this paper, we analyzed the effect of age on hyperalignment performance. Our findings demonstrate the significant impact of age-specific hyperalignment templates on the accuracy of individualized connectome prediction, neural response to the movie, as well as the inter-subject correlation (ISC) values after hyperalignment.

The ISC results demonstrate the age effect on hyperalignment template performance. Average ISC values in the hyperalignment common model space are higher when projecting individual connectome to the common model space derived from congruent age group templates—those aligned with the participant’s age group—rather than incongruent age group templates. By contrast, the congruency effect in the visual cortex is much smaller, suggesting that the aging effect on functional connectivity is smaller in these areas.

Moreover, congruent hyperalignment templates consistently achieved higher correlation between predicted individual connectomes and actual connectomes in participants’ native anatomical spaces compared to incongruent templates. When predicting an individual’s connectome in different age groups using either old group templates (> 65 years old) or young group templates (< 45 years old), as the individual’s age becomes more distant from the template age group, the prediction accuracy drops, and the difference between congruent and incongruent template prediction increases. Movie response prediction further indicates that the more congruent the template age group is with the participant’s age, the more accurate the prediction becomes.

We applied ISC analysis to an independent dataset, DLBS, to further validate the generalizability of our findings (see Supplementary Material). The results were consistent with those observed in the Cam-CAN dataset, showing that congruent templates yielded higher average ISC values than incongruent templates.

Previous hyperalignment study shows that the more data used in building the templates for estimating individual tuning matrices, the better the model performs (Feilong et al., 2023). We expect, therefore, that the effect of age-congruency of templates will be greater if the templates are built with larger data sets. Currently, the templates are built using only approximately 16 minutes of fMRI functional data from around 200 participants. We expect that better and more stable model performance could be achieved by incorporating data from more participants and increasing the amount of data per participant, preferably to at least an hour. Moreover, incorporating data using richer stimulus paradigms in template construction might provide additional information that could enhance template performance. Currently our templates are constructed using only resting state and sensorimotor task data. Naturalistic stimuli, such as movie - viewing, more broadly sample cognitive and brain states. With increased data collection and richer, naturalistic stimulus paradigms, we could capture a broader range of neural activities and idiosyncrasies, potentially leading to more robust and flexible hyperalignment templates and a more precise and reliable alignment between individual connectomes and the common model space.

In this work, we only focused on the impact of age on hyperalignment templates and observed a significant improvement using congruent age templates compared to incongruent ones. Based on these findings, it is important to consider the practical applications of this methodology in future research. When the goal of a case-control study is to directly compare functional organization or brain responses between distinct populations—such as clinical and non-clinical participants—it is essential that all individuals are hyperaligned to the same common template. For these analyses, researchers should either construct a joint template containing a balanced, representative sample from both groups, or align all participants to a normative control template based on age-matched data. This ensures that the resulting data share a single coordinate system, allowing for valid statistical comparisons between groups.

Age-specific or disease-specific templates are highly advantageous when the research objective is to maximize decoding accuracy or predictive performance within a specific population. Just as age introduces specific variations in neural architecture, neurological or psychiatric conditions—such as Alzheimer’s disease and Parkinson’s disease—can produce alterations in brain structure and function. In clinical or lifespan research, if the goal is to build a reliable biomarker of differences among individuals within a group, e.g., of disease progression or disease subtypes, or to map individualized connectomes for a specific patient cohort, researchers should use a template congruent with that specific group. By constructing disease-specific templates, it may be possible to capture these distinctive neural patterns with greater accuracy, leading to more precise alignment of individual connectomes. This approach could provide insights into the pathophysiological mechanisms underlying various clinical conditions and broaden the use of hyperalignment models in clinical neuroscience.

## References

Anderson, Z., Gratton, C., & Nusslock, R. (2021). The Value of Hyperalignment to Unpack Neural Heterogeneity in the Precision Psychiatry Movement. Biological Psychiatry: Cognitive Neuroscience and Neuroimaging, 6(9), 935–936. 10.1016/j.bpsc.2021.02.006

Anderson, Z., Turner, J. A., Ashar, Y. K., Calhoun, V. D., & Mittal, V. A. (2024). Application of hyperalignment to resting state data in individuals with psychosis reveals systematic changes in functional networks and identifies distinct clinical subgroups. Aperture Neuro, 4.10.52294/001c.91992

Cox, D. D., & Savoy, R. L. (2003). Functional magnetic resonance imaging (fMRI) “brain reading”:Detecting and classifying distributed patterns of fMRI activity in human visual cortex. NeuroImage, 19(2 Pt 1), 261–270. 10.1016/s1053-8119(03)00049-1

Esteban, O., Markiewicz, C. J., Blair, R. W., Moodie, C. A., Isik, A. I., Erramuzpe, A., Kent, J. D., Goncalves, M., DuPre, E., Snyder, M., Oya, H., Ghosh, S. S., Wright, J., Durnez, J., Poldrack, R. A., & Gorgolewski, K. J. (2019). fMRIPrep: A robust preprocessing pipeline for functional MRI. Nature Methods, 16(1), 111–116. 10.1038/s41592-018-0235-4

Feilong, M., Guntupalli, J. S., & Haxby, J. V. (2021). The neural basis of intelligence in fine-grained cortical topographies. eLife, 10, e64058. 10.7554/eLife.64058

Feilong, M., Jiahui, G., Gobbini, M. I., & Haxby, J. V. (2024). A cortical surface template for human neuroscience. Nature Methods, 21(9), 1736–1742. 10.1038/s41592-024-02346-y

Feilong, M., Nastase, S. A., Guntupalli, J. S., & Haxby, J. V. (2018). Reliable individual differences in fine-grained cortical functional architecture. NeuroImage, 183, 375–386. 10.1016/j.neuroimage.2018.08.029

Feilong, M., Nastase, S. A., Jiahui, G., Halchenko, Y. O., Gobbini, M. I., & Haxby, J. V. (2023). The individualized neural tuning model: Precise and generalizable cartography of functional architecture in individual brains. Imaging Neuroscience, 1, 1–34. 10.1162/imag_a_00032

Gratton, C., Laumann, T. O., Nielsen, A. N., Greene, D. J., Gordon, E. M., Gilmore, A. W., Nelson, S. M., Coalson, R. S., Snyder, A. Z., Schlaggar, B. L., Dosenbach, N. U. F., & Petersen, S. E. (2018). Functional Brain Networks Are Dominated by Stable Group and Individual Factors, Not Cognitive or Daily Variation. Neuron, 98(2), 439-452.e5. 10.1016/j.neuron.2018.03.035

Guntupalli, J. S., Feilong, M., & Haxby, J. V. (2018). A computational model of shared fine-scale structure in the human connectome. PLOS Computational Biology, 14(4), e1006120. 10.1371/journal.pcbi.1006120

Guntupalli, J. S., Hanke, M., Halchenko, Y. O., Connolly, A. C., Ramadge, P. J., & Haxby, J. V. (2016). A Model of Representational Spaces in Human Cortex. Cerebral Cortex (New York, N.Y.: 1991), 26(6), 2919–2934. 10.1093/cercor/bhw068

Haxby, J. V., Connolly, A. C., & Guntupalli, J. S. (2014). Decoding neural representational spaces using multivariate pattern analysis. Annual Review of Neuroscience, 37, 435–456. 10.1146/annurev-neuro-062012-170325

Haxby, J. V., Gobbini, M. I., Furey, M. L., Ishai, A., Schouten, J. L., & Pietrini, P. (2001). Distributed and Overlapping Representations of Faces and Objects in Ventral Temporal Cortex. Science, 293(5539), 2425–2430. 10.1126/science.1063736

Haxby, J. V., Guntupalli, J. S., Connolly, A. C., Halchenko, Y. O., Conroy, B. R., Gobbini, M. I., Hanke, M., & Ramadge, P. J. (2011). A common, high-dimensional model of the representational space in human ventral temporal cortex. Neuron, 72(2), 404–416. 10.1016/j.neuron.2011.08.026

Haxby, J. V., Guntupalli, J. S., Nastase, S. A., & Feilong, M. (2020). Hyperalignment: Modeling shared information encoded in idiosyncratic cortical topographies. eLife, 9, e56601. 10.7554/eLife.56601

Park, D. C., Hennessee, J. P., Smith, E. T., Chan, M. Y., Chen, X., Dakanali, M., Farrell, M. E., Liu, P.,Lu, H., Rofsky, N., Sun, X., Tamminga, C., Moore, W., Kennedy, K. M., Rodrigue, K., & Wig, G.S. (2025). The Dallas Lifespan Brain Study: A Comprehensive Adult Lifespan Data Set of Brain and Cognitive Aging. Scientific Data, 12(1), 846. 10.1038/s41597-025-04847-7

Taylor, J. R., Williams, N., Cusack, R., Auer, T., Shafto, M. A., Dixon, M., Tyler, L. K., Cam-CAN, & Henson, R. N. (2017). The Cambridge Centre for Ageing and Neuroscience (Cam-CAN) data repository: Structural and functional MRI, MEG, and cognitive data from a cross-sectional adult lifespan sample. NeuroImage, 144, 262–269. 10.1016/j.neuroimage.2015.09.018

