## Supplementary Materials for "Boosting Hyperalignment Performance with Age-specific Templates"

### Procedure building hyperalignment templates

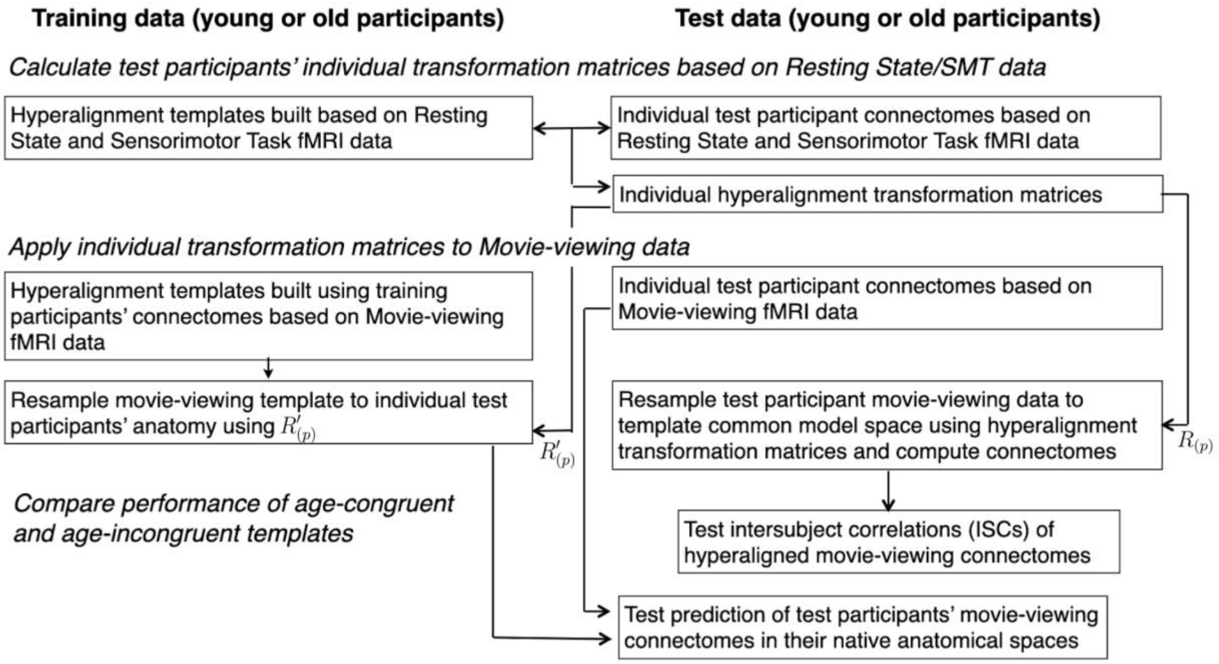


Supplementary Figure 1: Schematic of the procedure for building and testing hyperalignment templates

### Group differences in data quality

In the main analysis, we found that both ISC and prediction accuracy were higher for the young group than the old group, regardless of template type. We suspect that this might be related to data quality differences between the groups. Therefore, we performed an additional analysis, computing the temporal signal-to-noise ratio (tSNR) as a data quality measure, and we compared it between the two groups. We computed tSNR as the mean signal divided by the standard deviation for each participant and each vertex, using the resting-state data from the Cam-CAN dataset. We found that in general the average tSNR for the young group (mean ± standard deviation = 106.4 ± 21.2) is higher than the old group (93.9 ± 20.3), $t(429) = 6.25$, $p < 10^{-9}$.

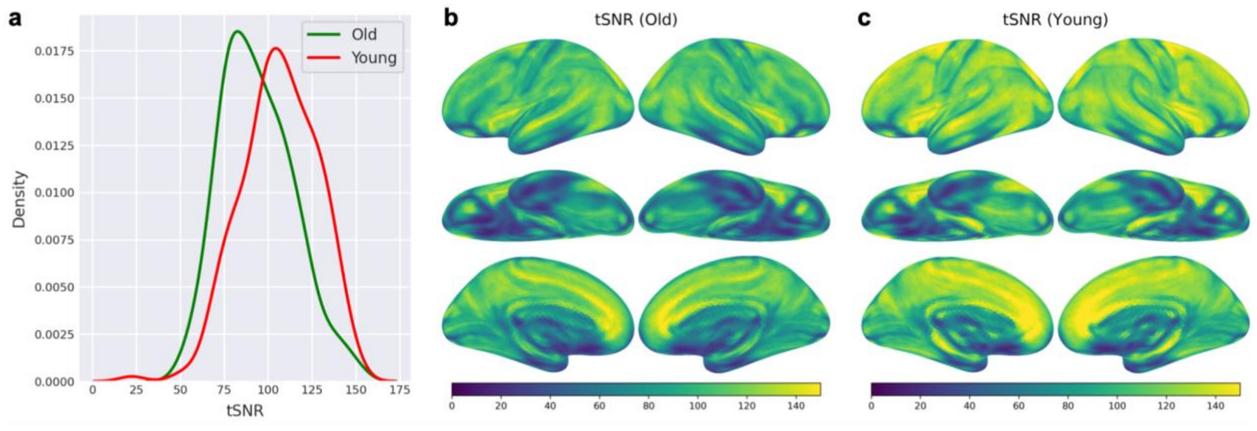


Supplementary Figure 2: tSNR differences between age groups. (**a**) The distribution of tSNR across participants, separately for each age group. (**b** and **c**) The distribution of tSNR across the cortex for the old group (**b**) and the young group (**c**).

### Validation Dataset – Dallas Lifespan Brain Study(DLBS)

The Dallas Lifespan Brain Study (DLBS) dataset collects multi-modal data from over 400 individuals across the entire adult lifespan, ranging from 20 to 90 years old. The dataset contains longitudinal data from multiple scan sessions conducted over 10 years. In this study, we focused on the fMRI data from the first and most complete scan session, which was collected between 2008 and 2014 from 464 participants. The fMRI data were collected using a Philips Achieva 3T scanner at the University of Texas Southwestern Medical Center, with 3.4 × 3.4 × 3.5 $mm^3$ voxels, TR = 2000 ms, TE = 25 ms, flip angle = 80° (Chan et al., 2014; Chen et al., 2021; Kennedy et al., 2015; Park et al., 2012). Among these participants, 407 completed a scanning session that included: 1) 7 minutes and 56 seconds of the words task, during which participants were shown 128 words and asked to make a semantic judgment on whether each word represented something living or nonliving; 2) three runs of 5 minutes and 56 seconds each of the scenes task, where participants were presented with outdoor landscape scenes and had to determine whether water was present in each scene; and 3) 6 minutes and 14 seconds of resting state. Among the 407 selected participants, 92 were younger than 45 years old, 133 were between 45 and 65 years old, and 182 were older than 65 years old.

The DLBS dataset was preprocessed using fMRIPrep version 24.1.0. Consistent with the procedure applied to the Cam-CAN dataset, we computed two types of functional connectomes based on different scan conditions:

1. using DLBS resting state and words task data,
2. using DLBS scenes task data.

### Inter-subject correlation

We applied the hyperalignment templates—constructed using Cam-CAN data from the two age groups (young and old)—to the DLBS dataset. Since the two datasets do not share identical functional tasks, we calculated transformation matrices that resample the resting state and words task fMRI data of the DLBS dataset into the Cam-CAN templates for young and old participants. We then applied the transformation matrices to connectomes based on the scenes task data and calculated the ISCs. The ISC results align well with those from the Cam-CAN dataset (Figure 3). We observed a statistically significant difference between ISC values derived from congruent and incongruent templates in both age groups (Supplementary Figure 3a). Although the differences among individual participants were small, they were still statistically significant with an anatomical distribution that is consistent with the congruency effects in the Cam-CAN dataset (Supplementary Figure 3b, 92.4% participants in the young group and 57.7% participants in the old group have higher congruent ISC values). SImilar to the results with the Cam-CAN data, we noted a stronger effect of template congruency on ISCs in the parietal, temporal, and occipital cortices, with smaller effects in occipital and ventral temporal visual cortices (Supplementary Figure 3c).

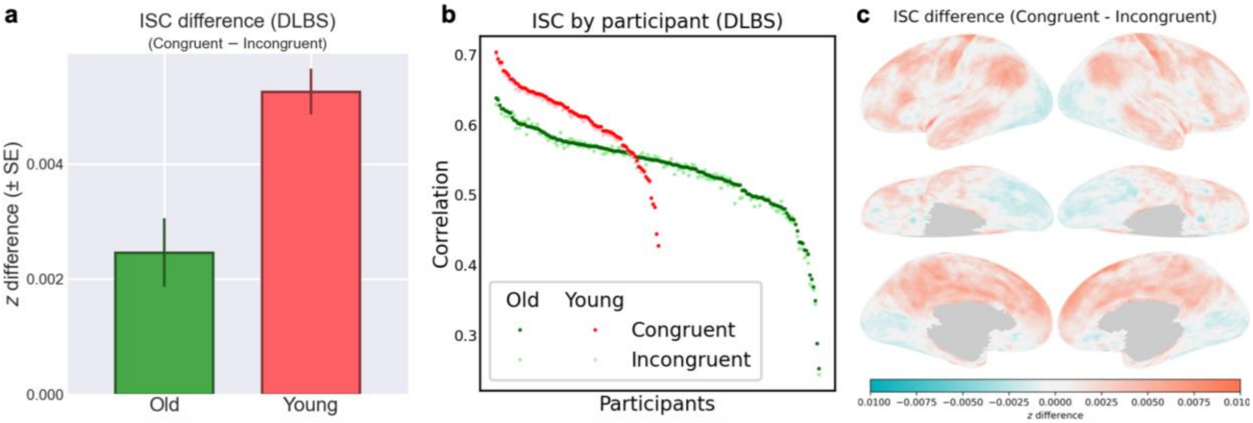


Supplementary Figure 3: Inter-subject Correlation Results (DLBS). (a) The average *z* (± SD) for the old group was 0.6044 ± 0.0789 for congruent templates, and 0.6020 ± 0.0810 for incongruent templates. The mean ISC difference (± SE) was 0.0025 ± 0.0006, $t(181) = 4.16$, Cohen's $d = 0.3087$ and $p < 10^{-4}$. The average *z* (± SD) for the young group was 0.7087 ± 0.0827 for congruent templates, and 0.7034 ± 0.0809 for incongruent templates. The mean ISC difference (± SE) was 0.0053 ± 0.0004, $t(91) = 13.24$, Cohen's $d = 1.3806$ and $p < 10^{-22}$. (b) Scatter plot of each participant’s average ISC value derived from different congruent and incongruent templates. (c) Topographic distribution of ISC difference across different brain regions

**ISC Cam-CAN middle-aged template**

We further evaluated the effect of age congruency by introducing an intermediate middle-aged cohort to our Cam-CAN dataset analysis. We computed the z-difference between ISC values after hyperalignment derived from congruent and incongruent templates to compare the middle-aged group against both the young and old cohorts (Supplementary Figure 4). When comparing the middle-age group to the young cohort, aligning middle-aged participants to their strictly age-congruent template actually resulted in slightly lower overall ISCs compared to using the young incongruent template (Supplementary Figure 4a). However, the young group maintained a strong congruency advantage over the middle-aged template. Conversely, when comparing the old cohort with the middle-aged cohort, both groups exhibited higher ISCs when aligned to their respective congruent templates (Supplementary Figure 4c). Breaking down the data by individual confirmed these distinct group-level trends across the majority of participants (Supplementary Figure 4b, d).

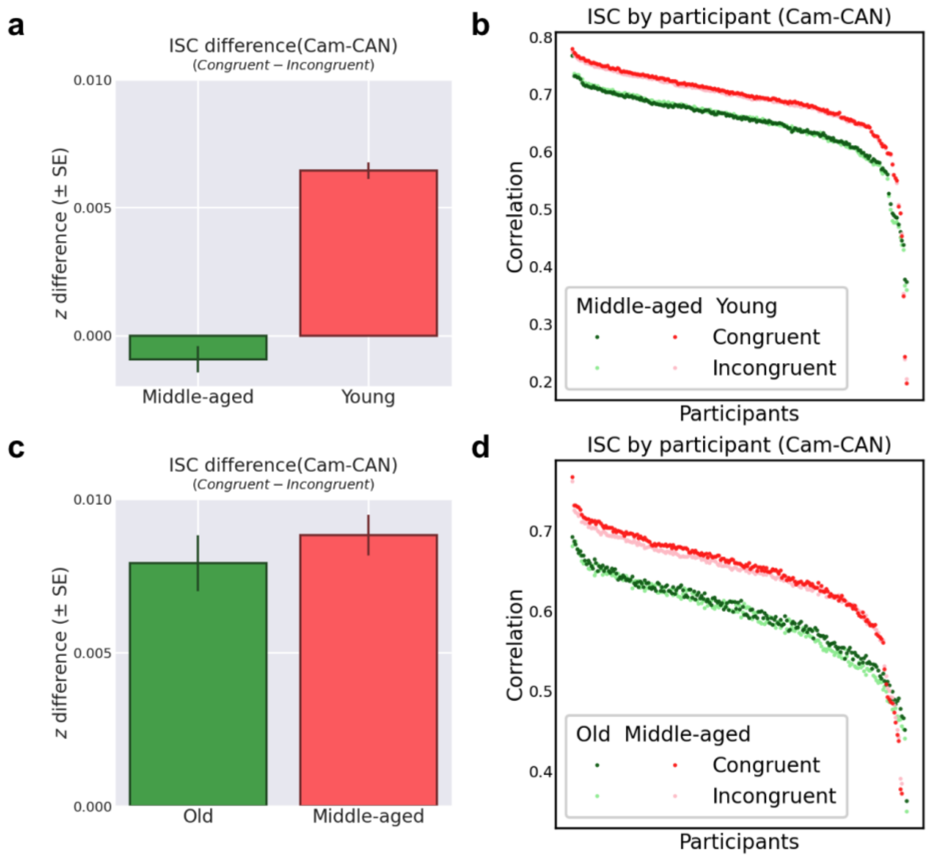


Supplementary Figure 4: Inter-subject Correlation Results based on the Middle-aged Cohort (Cam-CAN). (a) Mean ISC difference (Congruent – Incongruent) for the middle-aged and young groups. The mean ISC difference (± SE) for the middle-aged group was -0.0009 ± 0.0005, while the mean ISC difference for the young group was 0.0064 ± 0.0003. (b) Scatter plot of each participant's average ISC value derived from congruent and incongruent templates for the middle-aged versus young comparison. (c) Mean ISC difference (Congruent – Incongruent) for the old and middle-aged groups. The mean ISC difference (± SE) for the old group was 0.0079 ± 0.0009, and the mean ISC difference for the middle-aged group was 0.0088 ± 0.0007. (d) Scatter plot of each participant's average ISC value derived from congruent and incongruent templates for the old versus middle-aged comparison.

**Predicting connectomes Cam-CAN middle-aged template**

We also evaluated the impact of age congruency on fine-grained connectome prediction accuracy by incorporating the intermediate middle-aged cohort. We computed the z-difference between predicted and actual connectome correlations derived from congruent and incongruent templates. In contrast to the ISC results, connectome prediction demonstrated a consistent congruency advantage across all age comparisons. When comparing the middle-aged cohort to the young cohort, both groups exhibited significantly higher prediction accuracies when aligned to their respective age-congruent templates (Supplementary Figure 5a). Similarly, when comparing the old cohort with the middle-aged cohort, predictions generated using congruent templates were consistently more accurate than those using incongruent templates for both groups (Supplementary Figure 5c). Examining the breakdown by individual participant confirmed these robust group-level trends, illustrating that the vast majority of participants across all age groups achieved optimal connectome prediction when aligned to a template congruent with their specific age cohort (Supplementary Figure 5b, d).

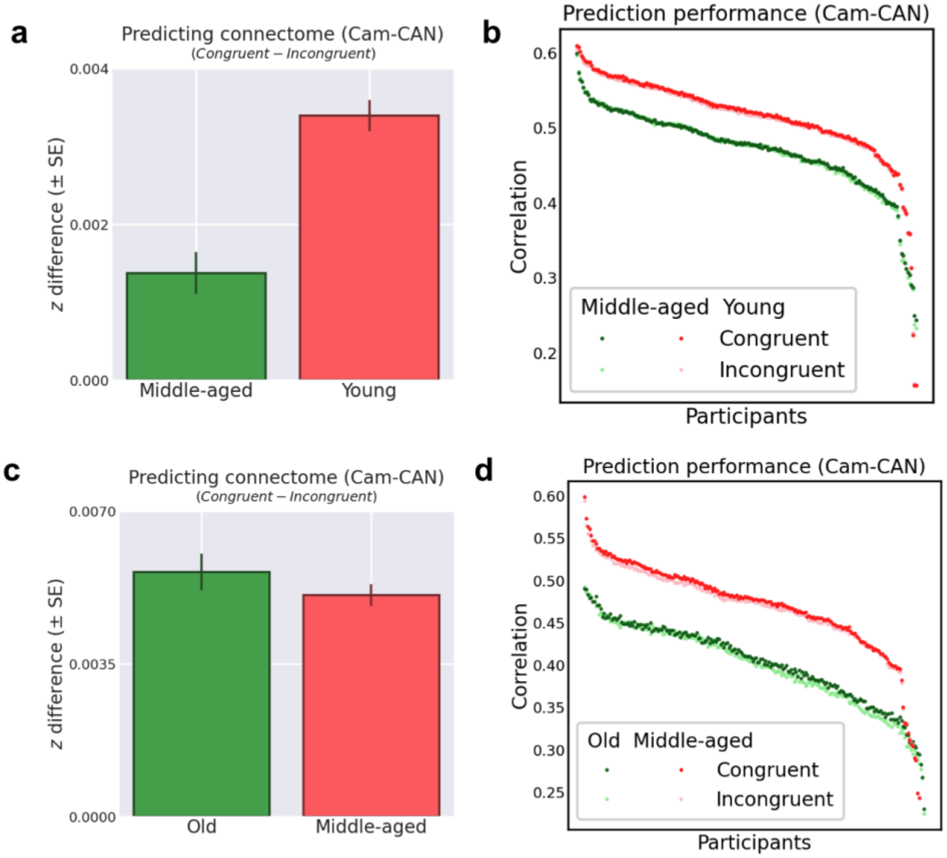


Supplementary Figure 5: Connectome Prediction Results based on the Middle-aged Cohort (Cam-CAN). (a) Mean z-difference (Congruent – Incongruent) in connectome prediction accuracy for the middle-aged and young groups. The mean difference (± SE) for the middle-aged group was 0.0014 ± 0.0003, while the mean difference for the young group was 0.0034 ± 0.0002. (b) Scatter plot of each participant's connectome prediction correlation derived from congruent and incongruent templates for the middle-aged versus young comparison. (c) Mean z-difference (Congruent – Incongruent) in connectome prediction accuracy for the old and middle-aged groups. The mean difference (± SE) for the old group was 0.0056 ± 0.0004, and the mean difference for the middle-aged group was 0.0051 ± 0.0002. (d) Scatter plot of each participant's connectome prediction correlation derived from congruent and incongruent templates for the old versus middle-aged comparison.

**Predicting connectomes Cam-CAN with 10-year age templates**

We have constructed templates using narrower, 10-year age intervals and evaluated their performance. Because different age groups have different numbers of people, we use a fixed number of training participants for each age group (two thirds of the people from the group with the minimal number of people) to build the templates to make a fair comparison. The results show a continuous gradient of age-related divergence. When predicting data for the 80–90 cohort, the 20–30 template performs the worst and the performance steadily improves as the template age gets closer to the target demographic. This systematic gradient further supports our main finding: the penalty for using an incongruent template increases with the discrepancy between the template age and participant age.

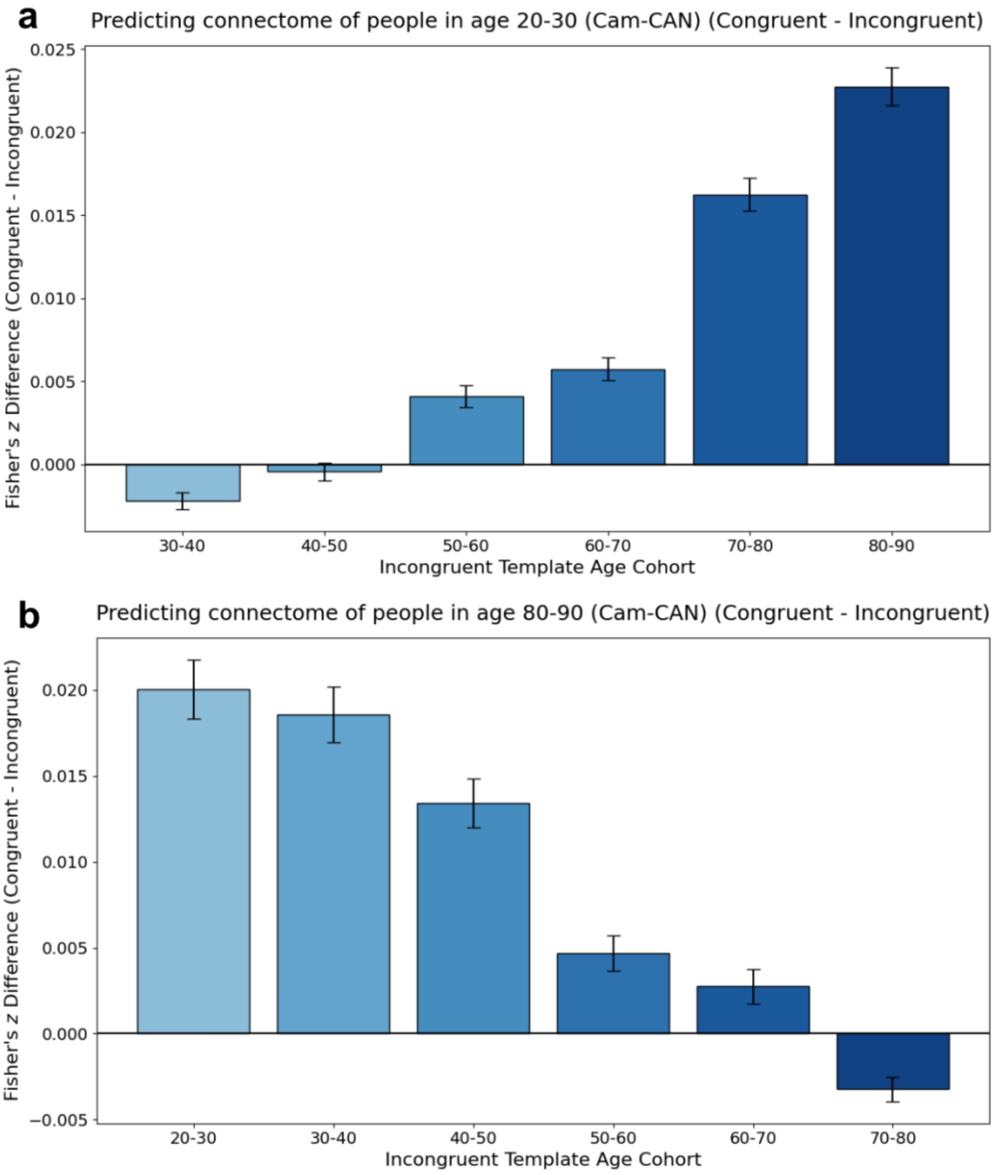


Supplementary Figure 6: Connectome Prediction Results with 10-year age templates: **(a)** Mean Fisher's z difference (Congruent – Incongruent) in connectome prediction for the young cohort (20–30 years old). The bar plot compares the performance of the age-congruent template against incongruent templates from progressively older cohorts (ranging from 30–40 to 80–90). **(b)** Mean Fisher's z difference (Congruent – Incongruent) in connectome prediction for the older cohort (80–90 years old), comparing the age-congruent template against incongruent templates from progressively younger cohorts (ranging from 20–30 to 70–80).
